# Legionella uses host Rab GTPases and BAP31 to create a unique ER niche

**DOI:** 10.1101/2024.05.10.593622

**Authors:** Attinder Chadha, Yu Yanai, Hiromu Oide, Yuichi Wakana, Hiroki Inoue, Saradindu Saha, Mitsuo Tagaya, Kohei Arasaki, Shaeri Mukherjee

**Affiliations:** G.W. Hooper Foundation, University of California at San Francisco, San Francisco, USA; Department of Microbiology and Immunology, University of California at San Francisco, San Francisco, USA; Chan Zuckerberg Biohub, San Francisco, USA; School of Life Sciences, Tokyo University of Pharmacy and Life Sciences, Hachioji, Tokyo 192-0392, Japan

**Author notes:** Authors contributed equally (Attinder Chadha, Kohei Arasaki).

## Abstract

Upon entry into host cells, the facultative intracellular bacterium *Legionella pneumophila* (*L.p*.) uses its type IV secretion system, Dot/Icm, to secrete ~330 bacterial effector proteins into the host cell. Some of these effectors hijack endoplasmic reticulum (ER)-derived vesicles to form the *Legionella*-containing vacuole (LCV). Despite extensive investigation over decades, the fundamental question persists: Is the LCV membrane distinct from or contiguous with the host ER network? Here, we employ advanced photobleaching techniques, revealing a temporal acquisition of both smooth and rough ER (sER and rER) markers on the LCV. In the early stages of infection, the sER intimately associates with the LCV. Remarkably, as the infection progresses, the LCV evolves into a distinct niche comprising an rER membrane that is independent of the host ER network. We discover that the *L.p.* effector LidA binds to and recruits two host proteins of the Rab superfamily, Rab10, and Rab4, that play significant roles in acquiring sER and rER membranes, respectively. Additionally, we identify the pivotal role of a host ER-resident protein, BAP31, in orchestrating the transition from sER to rER. While previously recognized for shuttling between sER and rER, we demonstrate BAP31’s role as a Rab effector, mediating communication between these ER sub-compartments. Furthermore, using genomic deletion strains, we uncover a novel *L.p.* effector, Lpg1152, essential for recruiting BAP31 to the LCV and facilitating its transition from sER to rER. Depletion of BAP31 or infection with an isogenic *L.p.* strain lacking Lpg1152 results in a growth defect. Collectively, our findings illuminate the intricate interplay between molecular players from both host and pathogen, elucidating how *L.p.* orchestrates the transformation of its residing vacuole membrane from a host-associated sER to a distinct rER membrane that is not contiguous with the host ER network.

## INTRODUCTION

The bacterium *Legionella pneumophila* (*L.p.)* is the causative agent of a severe type of pneumonia known as Legionnaires’ disease. *L.p.* primarily affects immunocompromised and elderly patients, and infection can often lead to a high mortality rate^1^. When aerosols containing *L.p.* are generated, they can enter the human lung, where it is detected and phagocytosed by alveolar macrophages. Upon entry, *L.p.* uses its type IV secretion system, Dot/Icm, to secrete more than 300 bacterial effector proteins into the host cell^2,3^. Despite their identification, the targets and functions of most of these *L.p.* effectors are currently unknown^4^. For the effectors whose functions are known, *L.p.* has been shown to target host proteins that are evolutionarily conserved from amoeba to humans, many of which are key players in regulating membrane traffic in cells^5–9^. Studying the function of *L.p.* effectors has often led to a deeper understanding of the role of essential host proteins, including a novel posttranslational modification^5^ and non-canonical ubiquitination signaling^10–14^. Some of these effectors are thought to play key roles in acquiring host endoplasmic reticulum (ER)-derived vesicles to form the *Legionella*-containing vacuole (LCV)^15^. Improper ER membrane recruitment leads to defects in bacterial replication^16,17^. However, two big questions remain: 1) Is the LCV contiguous with the host ER membrane, or is it a separate entity? 2) What effectors and host proteins help create this unique LCV niche?

The ER is organized into a ribosome binding compartment, termed rough ER (rER), and a non-ribosome binding compartment, termed the smooth ER (sER)^18^. The sER has a tubular structure and functions in Ca^2+^ signaling and lipid synthesis. Its structure is maintained by a wide range of membrane-shaping proteins, including Reticulon-4 (Rtn4)^19^. In contrast, the rER has a sheet-like structure and functions in *de novo* protein synthesis^18^. The protein Sec61β functions as a translocon that facilitates protein entry into the ER and is associated with the rER^20^. The composition of the ER membrane, however, is dynamic. This is best characterized by ER resident proteins that shuttle between sER and rER. One such protein is B-cell receptor-associated protein 31 (BAP31)^21^. BAP31 is a transmembrane protein that primarily localizes to the sER under steady-state conditions but translocates to the rER under conditions when the ER is perturbed^22^. Due to the continuous exchange of membranes within the ER, the ratio of the rER to the sER in cells is not always constant. This variability in ER sub-compartment composition influences various cellular functions, such as secretory capacity and translation ability^23^. By creating an ER-derived niche, *L.p*. provides an opportunity to explore the intricate communication between different ER sub-compartments.

In this study, we aimed to leverage *L.p.* as a discovery tool to uncover the communication dynamics between ER sub-compartments. Previous observations indicated ribosome presence on the *Legionella*-containing vacuole (LCV) surface, suggesting a rough ER (rER) association^24^. However, recent data shows that sER markers such as Rtn4 are ubiquitinated and recruited to the LCV^25^. Here, we try to resolve this seemingly disparate observation by providing the first evidence that the LCV transitions from a sER association to a rER identity in a temporal manner. Interestingly, it can maintain a unique rER niche that is not contiguous with the host ER membrane. *L.p.* utilizes its meta effector LidA to acquire host Rab GTPases, Rab10 and Rab4, on the LCV. These GTPases and the ER-resident protein, BAP31, allow *L.p.* to mature from a sER to a rER identity. Notably, we identify Lpg1152 as a key *L.p.* effector that binds BAP31, facilitating LCV maturation during infection. Thus, our study reveals BAP31 as a novel Rab effector crucial for ER sub-compartment communication, underscoring the elaborate mechanisms by which *L.p.* manipulates host proteins and effectors to rewire the ER network.

## Result

### *L.p.* associates temporally with the sER and later creates a unique rER niche independent of the host ER network

Given that *L.p.* has been shown to associate with both sER and rER membranes^24,25^, we sought to understand whether acquiring these markers was a temporally regulated process. We used a HeLa cell line that stably expresses FcγRII (HeLa Fcγ) to facilitate *L.p.* entry via antibody-mediated opsonization. We analyzed the sER *vs*. rER association of *L.p.* by confocal microscopy and used Rtn4 and Sec61β as sER and rER markers, respectively. Interestingly, we observed a remarkable temporal segregation of sER and rER markers on the LCV. At the initial stage of infection (2 hours), the LCV showed an enrichment of Rtn4 (**Fig. 1A top panel**), suggesting a sER localization. In contrast, at a late stage (8 hours), the LCV showed rER association, as evident by Sec61β staining (**Fig. 1A bottom panel**). This data is consistent with previous EM and confocal data that have demonstrated the enrichment of these markers on the LCV^25–27^. Next, we sought to answer a long-standing question in the field: is the LCV membrane separate or contiguous with the host ER membrane? We chose two independent photobleaching techniques to answer this question. First, we employed fluorescence loss in photobleaching (FLIP) time-lapse imaging. Briefly, FLIP is used to assess the continuity between areas within cells and involves the repeated photobleaching of a small region of a cell that expresses a fluorescent marker. Over time, because of the lateral movement of fluorophores, fluorescence loss is seen in other regions of the cell that are interconnected with the bleached area. In contrast, the fluorescence in the unconnected areas is unaffected because fluorescent markers do not pass through the bleached region. We utilized HeLa Fcγ cells that stably express a sER marker (GFP-Rtn4) and a rER marker (RFP-Sec61β). At 3 hours post-infection, continuous photobleaching of Rtn4 in a region in the ER (red box) which is distant from the LCV membrane (purple box) caused a concomitant loss of the Rtn4 signal both in the host ER network as well as the LCV membrane (**Fig. 1B and 1C; Movie S1A**), suggesting that the LCV membrane is contiguous with the host sER network. However, at 6 hours post-infection, even though the photobleaching of Sec61β in ER regions (red box) distant from the LCV (purple box) results in complete loss of fluorescence in the entire host ER membrane, Sec61β fluorescence on the LCV remained completely protected (**Fig. 1D and 1E; Movie S1B and S1C**). This striking result indicates that the ER membrane on the LCV at 6 hours post-infection is a distinct entity entirely independent of the host ER membrane network. To confirm these findings, we subjected HeLa Fcγ cells to fluorescence recovery after photobleaching (FRAP). FRAP involves the photobleaching of fluorescently tagged proteins via brief, intensive laser excitation followed by measuring the fluorescence recovery rates of the fluorophores back into the briefly bleached area. If the fluorophores are in distinct membrane compartments, there is little recovery. However, interconnected membranes will show recovery of fluorescence over time. After 6 hours post-infection, photobleaching of Sec61β on the LCV (red box) leads to a significant reduction in fluorescence intensity at the LCV (white arrow) compared to the distal ER (blue box) (**Fig. 1F and 1G; Movie S1D and S1E**). By 40 seconds post bleaching, the host ER (blue box) recovers ~80% of the Sec61β signal, whereas the LCV (red box) only recovers ~20%. This confirms our FLIP findings that, at this late stage, the LCV has a distinct rER membrane separate from the host ER. Taken together, the FLIP and FRAP results unequivocally show that the LCV moves from a host-associated continuous sER membrane to a distinct rER membrane that is independent of the host ER network.

**Figure 1.**
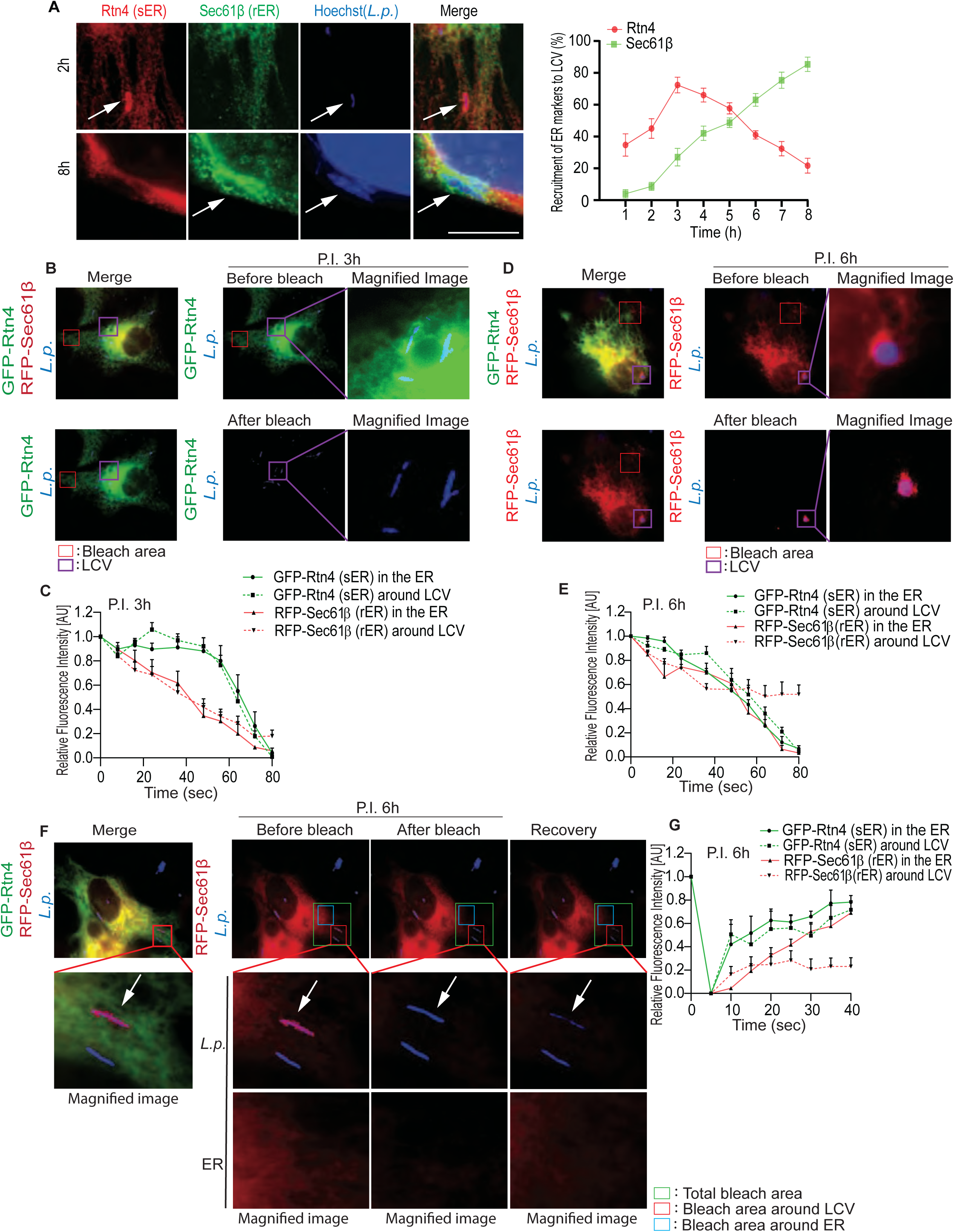
*L.p.* temporally associates with the sER and rER and later creates a unique ER niche independent of the host ER network. (A) HeLa Fcγ cells were infected with *L.p.* for the indicated times, stained with anti-Rtn4 (sER marker), Sec61β (rER marker) antibodies, and Hoechst (*L.p*.) for immunofluorescence analysis, bar = 5 µm. Results represent three independent experiments (100 vacuoles were scored in each). Error bars represent mean ± SD. (B) HeLa Fcγ cells stably expressing GFP-Rtn4 and RFP-Sec61β were infected with Halo-tagged WT *L.p*. for 3 hours for FLIP. The top panel represents GFP-Rtn4 before photobleaching, and the bottom represents GFP-Rtn4 after photobleaching for 80 seconds. (C) The graph represents the relative fluorescence intensity of GFP-Rtn4 (green lines) and RFP-Sec61β (red lines) in the ER and around LCV. Error bars represent mean ± SD. The purple and red boxes represent the area around LCV and bleach areas, respectively. (D) HeLa Fcγ cells stably expressing GFP-Rtn4 and RFP-Sec61β were infected with WT *L.p*. for 6 hours. The top panel represents RFP-Sec61β before photobleaching, and the bottom represents RFP-Sec61β after photobleaching. The purple and red boxes represent the area around the LCV and bleach areas, respectively. (E) The graph represents the relative fluorescence intensity of GFP-Rtn4 (green lines) and RFP-Sec61β (red lines) in the ER and around LCV. (F) HeLa Fcγ cells stably expressing GFP-Rtn4 and RFP-Sec61β were infected for 6 hours, and RFP-Sec61β was photobleached for FRAP. The green box represents the total area of bleach. The red and blue boxes represent the bleached area around the LCV and the ER, respectively. White arrows represent the area around LCV. (G) The graph represents the relative fluorescence intensity of GFP-Rtn4 (green lines) and RFP-Sec61β (red lines) in the ER and around LCV. Error bars represent mean ± SD.

### Rab4 and Rab10 facilitate the transition of the LCV from a sER to a rER identity

Our previous results stimulated our interest in identifying molecular mechanisms that facilitate LCV maturation from the sER to the rER. Rab GTPases (members of the Ras superfamily) regulate the maturation of the endosomal and secretory pathway compartments^28^. Indeed, several *L.p.* effectors are known to manipulate and modify these Rab GTPases^29^. One such effector, LidA, was shown to interact with Rab1^30^. Subsequent work described LidA as a Rab GTPase supereffector that forms a stable stoichiometric complex with at least two other Rabs, Rab6 and Rab8a^31^. Despite this, the entire repertoire of interacting partners of LidA has not been identified. Upon entry, *L.p.* is known to first interact with the endosomal compartment, followed by the acquisition of host ER vesicles^15^. This observation prompted us to ask if early endocytic Rabs, such as Rab4 or Rab5, and Rab10, an ER-specific Rab that regulates ER morphology and dynamics^32^ and is required for optimal *L.p.* growth^33^, interact with LidA.

To investigate this, HEK293 Fcγ cells were transfected with either 3X-FLAG-Rab4, 3X-FLAG-Rab5, or 3x-FLAG-Rab10 and LidA. Surprisingly, LidA preferentially interacted with Rab4 and Rab10 but not with Rab5 (**Fig. 2A**). These results suggested that LidA could indeed act as a Rab super effector that allows the LCV to temporally interact with the early endocytic compartment followed by the ER network.

**Figure 2.**
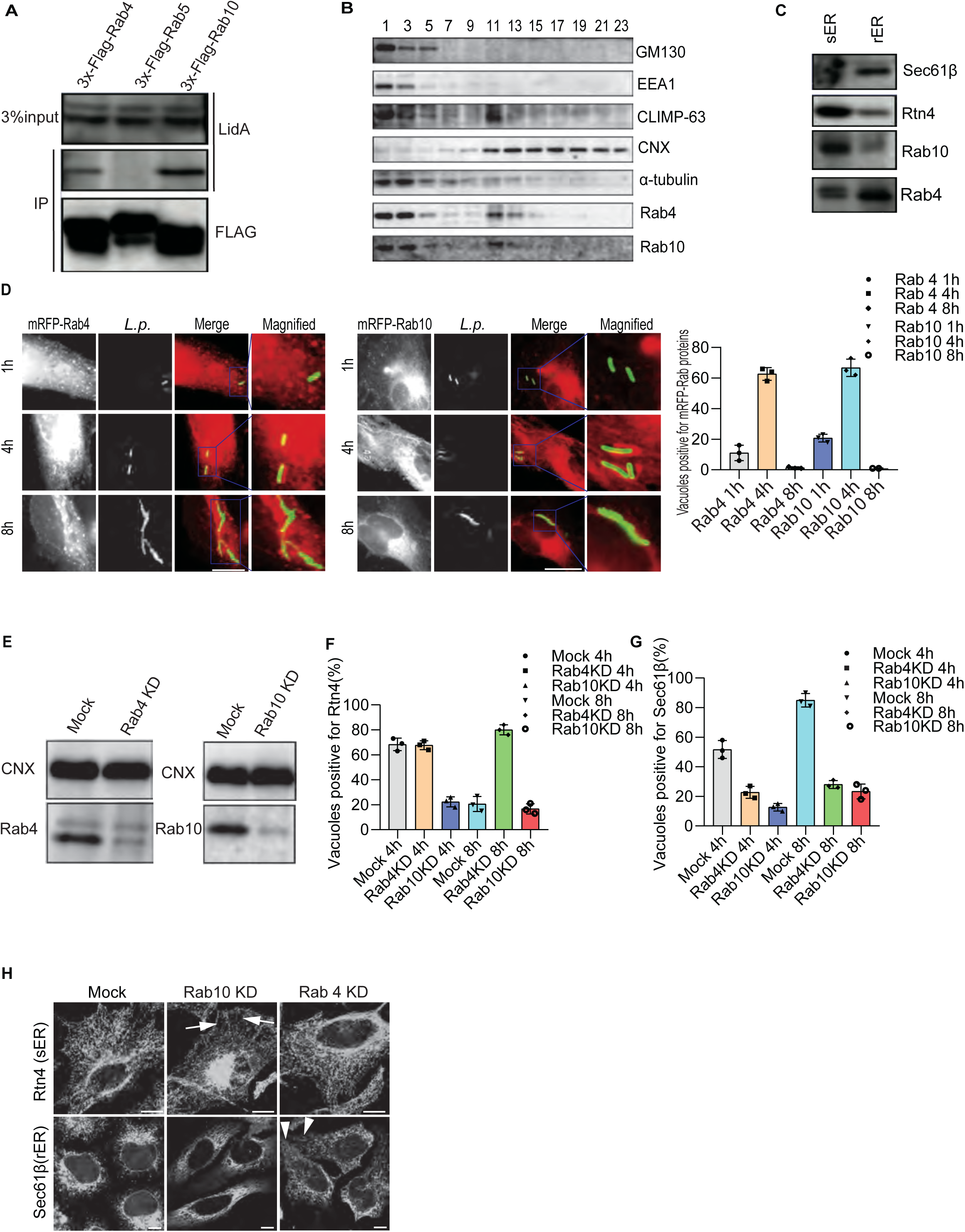
Rab4 and Rab10 facilitate the transition of the LCV from a sER to a rER identity. (A) HEK293-Fcγ cells were transfected with the indicated plasmids. After transfection, cell lysates were immunoprecipitated with anti-FLAG antibody and subjected to immunoblot analysis against the indicated antibodies. 3% input and immunoprecipitated proteins are indicated. (B) HeLa-Fcγ cells were subjected to subcellular fractionation, and immunoblot analysis was done against the indicated proteins. (C) Sucrose gradient fractionation was done to obtain the sER and rER fractions and immunoblotted against indicated antibodies. (D) HeLa-Fcγ cells expressing mRFP-Rab4 or mRFP-Rab10 were infected with *L.p.* for the indicated times, stained with *L.p*. for immunofluorescence analysis, and the vacuoles positive for mRFP-Rab4 or -Rab10 were quantified. Bar = 5 µm. The purple box represents the magnified image. Results represent three independent experiments (100 vacuoles were scored in each). Error bars represent mean ± SD. (E) HEK293-Fcγ cells were transfected with either mock, Rab4, or Rab10 siRNA for 72 hours, and cell extracts were immunoblotted against the indicated antibodies. (F, G) HeLa Fcγ cells were silenced for Rab4 or Rab10 for 72 hours, infected with *L.p.* for 4 and 8 hours, and the vacuoles positive for Rtn4 or Sec61β were counted. Results represent three independent experiments, and statistical significance was measured using one-way ANOVA. (100 vacuoles were scored in each experiment). Bars represent mean ± SD. (H) HeLa Fcγ cells were silenced with Rab4 and Rab10 siRNA, and the ER morphology was observed by immunofluorescence analysis. Bars = 5 µm. Arrows indicate disruption of the sER tubular network. Arrowheads indicate diffusion of rER to the cell periphery.

Guided by these findings, we next interrogated the recruitment of Rab4 and Rab10 on the LCV. To assess this, we first confirmed the presence of Rab4 and Rab10 in the ER by subcellular fractionation (**Fig. 2B**). ER-derived membrane fractions (fractions 11-23) were separated from Golgi- and endosome-derived fractions (fractions 1-5). Even though Rab4 and Rab10 were detected in the endosomal fractions (1-3), we also detected a significant amount of Rab4 and Rab10 in fractions containing ER-derived membrane (fractions 11-19) (**Fig. 2B**). Moreover, the separation of rER and sER membranes from whole cell membranes of HEK293-Fcγ cells revealed that Rab10 was predominantly enriched in Rtn4-positive sER whereas Rab4 was abundant in Sec61β-positive rER (**Fig. 2C**). Indeed, confocal microscopy after 4 hours post-infection with WT *L.p*. revealed that both Rab4 and Rab10 were associated with the LCV membrane (**Fig. 2D**), suggesting a role for these small GTPases in the LCV maturation process. To test this notion, we silenced Rab4 and Rab10 in Hela Fcγ cells with small interfering RNAs (siRNAs) (**Fig. 2E),** infected these cells with *L.p.,* and assayed for the association of Rtn4 and Sec61β with the LCV membrane (**Fig. 2F** and **Fig. 2G**). Interestingly, the LCV in Rab10 silenced cells showed dramatic reduction in acquiring either Rtn4 (after 4 hours) or Sec61β (after 8 hours) (**Fig. 2F** and **Fig. 2G**). In contrast, LCVs in Rab4 silenced cells showed no change in Rtn4 acquisition after 4 hours. However, 8 hours post-infection, Rab4 silencing significantly increased Rtn4 positive vacuoles compared to mock treatment and showed a defect in acquiring Sec61β (**Fig. 2F and Fig. 2G**). These results suggested that Rab10 functions upstream of Rab4 during the LCV maturation process and that LCV maturation from sER to rER is at least partly dependent on Rab4 (**Fig. 2F and 2G**). In contrast, recruitment of Sec61β on LCV is downstream of both Rab4 and −10, as silencing of both Rab4 and −10 significantly reduced the vacuoles that were positive for Sec61β at 8 hours post-infection (**Fig. 2F and 2G**). It is also possible that Rab4 is required to remove sER, which sets the stage for rER acquisition.

Given that both Rab4 and −10 are abundant in rER and needed for LCV transition from sER to rER, we next asked if Rab10 and Rab4 are essential for maintaining the ER sub-compartment architecture. To demonstrate this, Rab10 and Rab4 were silenced in HeLa Fcγ cells and then stained for Rtn4 and Sec61β. Interestingly, silencing of Rab10 disrupted sER morphology (white arrows, **Fig. 2H**), as evident from the less pronounced meshwork of Rtn4 staining in the periphery. Upon Rab4 knockdown, the rER staining (Sec61β) was reduced in the reticular region and appeared to relocate to the periphery, suggesting distorted rER (white solid arrows, **Fig. 2H**). These data point to the crucial role these small GTPase molecules play in maintaining the ER dynamics and morphology. Taken together, these results implicate an interesting temporal kinetics of recruitment of the s-ER marker on LCV early during infection, followed by the r-ER. These findings also indicate the involvement of Rab4 and Rab10 during LCV maturation and their role in maintaining ER sub-compartment morphology.

### BAP31 binds to Rab4 and Rab10 while BAP29 interacts with Rab4

Rab proteins act as molecular switches on the membrane and carry out their function with the help of other proteins known as Rab effectors^28^. We reasoned that a Rab effector protein may bind to Rab4 and Rab10 to help facilitate LCV maturation from the sER to the rER. Previous work identified BAP31 as an ER resident protein that shuttles between the two ER sub-compartments^22^. Furthermore, it was shown to bind to Rab11a, a Rab family member that regulates endocytic recycling^34^. Interestingly, its homolog BAP29, which has ~50% sequence identity with BAP31, resides in the rER and does not shuttle to sER^35^. Together, these data suggest that BAP31 could be a strong candidate to serve as a Rab4/Rab10 effector protein and facilitate LCV maturation.

First, we performed sucrose gradient fractionation to confirm the localization of BAP31 and BAP29 to ER sub-compartments. As reported previously, BAP31 was detected equally in both sER and rER fractions. In contrast, BAP29 was predominantly localized to the rER, with a small fraction also present in the sER (**Fig. 3A**). To demonstrate the *in vivo* interaction of Rab4/10 with BAP31/29, we performed proximity ligand assay (PLA). Proximity ligand assay (PLA) is used to identify *in vivo* interactions between two proteins within 30-40 nm distance. In this assay, cells are fixed and probed with primary antibodies against the proteins of interest. Secondary antibodies with PLA oligonucleotide probes are added that complement each other. A polymerase then amplifies the signal, and fluorescently labeled oligonucleotide probes bind to the amplified DNA. Each signal comprises ~1000 bound fluorescent probes that appear as punctate dots under a microscope. Interestingly, our results show that BAP31 associates with both Rab4 and Rab10 (**Fig. 3B and 3D**), which may partly explain the shuttling of BAP31 between the sER and the rER. Consistent with the localization in the ER subdomain, BAP29 preferentially interacts with Rab4 and not Rab10 (**Fig. 3C and 3E**). The percentage of co-localization has been presented as a histogram (**Fig. 3F**). To further validate these findings, an immunoprecipitation assay was performed. Briefly, HEK293 Fcγ cells were transfected with either 3x-FLAG empty vector (negative control), 3x-FLAG-Rab4, or 3x-FLAG-Rab10 plasmids. Post transfection, cell lysates were immunoprecipitated with a FLAG antibody, and eluates were subjected to SDS-PAGE and immunoblotted with the antibodies against BAP31, FLAG, and BAP29. Consistent with the PLA results, BAP31 immunoprecipitated with both 3x-FLAG-Rab10 and 3x-FLAG-Rab4, whereas BAP29 was mostly immunoprecipitated with 3x-FLAG-Rab4 and very little with 3x-FLAG-Rab10 (**Fig. 3G and 3H**). Collectively, our results show that BAP31 or BAP29 are novel interacting partners of Rab4 and Rab10 and perhaps are required for intercompartment shuttling in the ER.

**Figure 3.**
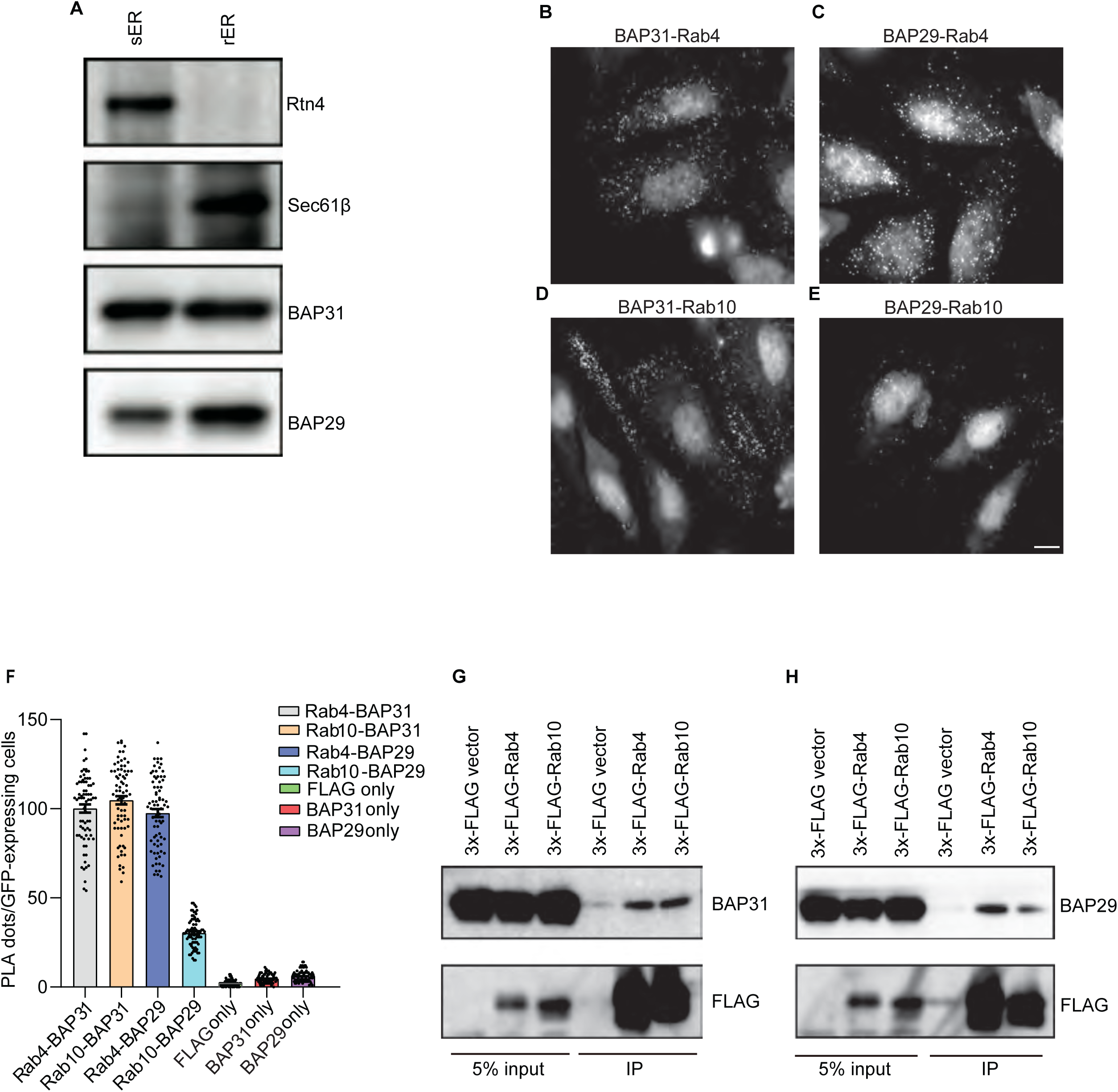
BAP31 binds to Rab4 and Rab10, while BAP29 interacts with Rab4. (A) The membranes of sER and rER fractions were collected by centrifugation and subjected to immunoblotting against the indicated antibodies. (B-E) HeLa-Fcγ cells expressing GFP vector, 3x-FLAG-Rab4 or 3XFLAG-Rab10 were fixed, PLA (Proximity ligation assay) was performed using the indicated antibodies, and the number of PLA dots was quantified. Bar = 5 µm. (F) Results were representative of three independent experiments, and statistical significance was achieved using one-way ANOVA. (PLA dots in 25 cells expressing GFP were scored in each experiment). Error bars represent mean ± SD. (G, H) HEK293-Fcγ cells were transfected with the indicated plasmids, and cell extracts were subjected to immunoprecipitation assay using the anti-FLAG antibody and immunoblotted against the indicated antibodies.

### BAP31 is required for the sER to rER transition and efficient replication of *Legionella*

Due to the interaction of BAP31 with both Rab10 and Rab4, pivotal players in LCV membrane maturation, we aimed to ascertain *L.p.*’s reliance on BAP31 to facilitate this crucial process. The initial examination focused on visualizing the recruitment of BAP31 to the LCV membrane. We noted robust recruitment of BAP31 (shown in green within the white box) on the LCV, peaking at 4 hours post-infection and persisting throughout the 8-hour duration (**Fig. 4A**). Subsequently, we hypothesized that BAP31 might be integral to LCV maturation. To explore this hypothesis, we silenced BAP31 in HeLa Fcγ cells using siRNAs (**Fig. 4B**) and infected them with WT *L.p.* for specified durations, followed by immunostaining for Rtn4 (red) and Sec61β (green). Consistent with prior observations, infection of HeLa Fcγ cells treated with mock siRNA demonstrated temporal recruitment of ER markers to the LCV, with Rtn4 peaking at 3 hours and Sec61β at 7 hours (**Fig. 4C**). In contrast, silencing BAP31 led to an enrichment of Rtn4 (green) on the LCV, accompanied by a depletion of Sec61β (red). These findings suggest a crucial role for BAP31 in orchestrating LCV maturation from the sER to the rER (**Fig. 4D and 4E**). To test if preventing LCV membrane maturation affects *L.p.* replication, we silenced BAP31 and conducted a replicative vacuole assay. BAP31 depletion resulted in a marked reduction in the number of *L.p*. in the LCV (8-hour post-infection) (**Fig. 4F**). Next, we probed if recruitment of Rab4/10 and BAP31/29 was dependent on each other during LCV maturation. To do this, first, we checked the recruitment of BAP31 (green) and BAP29 (green) in Rab4 and Rab10 depleted cells. Intriguingly, the silencing of Rab10 impaired the recruitment of BAP31, compared to mock and Rab4 siRNA (**Fig. S2A and B**). In contrast, the recruitment of BAP29 was abrogated from both Rab4/10 silenced cells (**Fig. S2 C and D**). These data further establish the idea that BAP proteins are Rab effectors and are recruited to membranes in a Rab-dependent manner. Next, we explored the dependence of Rab4/Rab10 recruitment to LCVs on BAP31/BAP29 (**Fig. 4B and Fig. S3A)**. As expected, Rab10 (red) recruitment on the LCVs was independent of BAP31 and BAP29 **(Fig. S3 B and C)**. However, the recruitment of Rab4 (red) was strongly dependent on BAP31 since silencing of BAP31 caused a dramatic reduction in the mRFP-Rab4 positive vacuoles **(Fig. S3 D and E)**. These results suggest Rab10 and BAP31 play an essential role upstream in recruiting Rab4 and other rER markers to the LCV.

**Figure 4.**
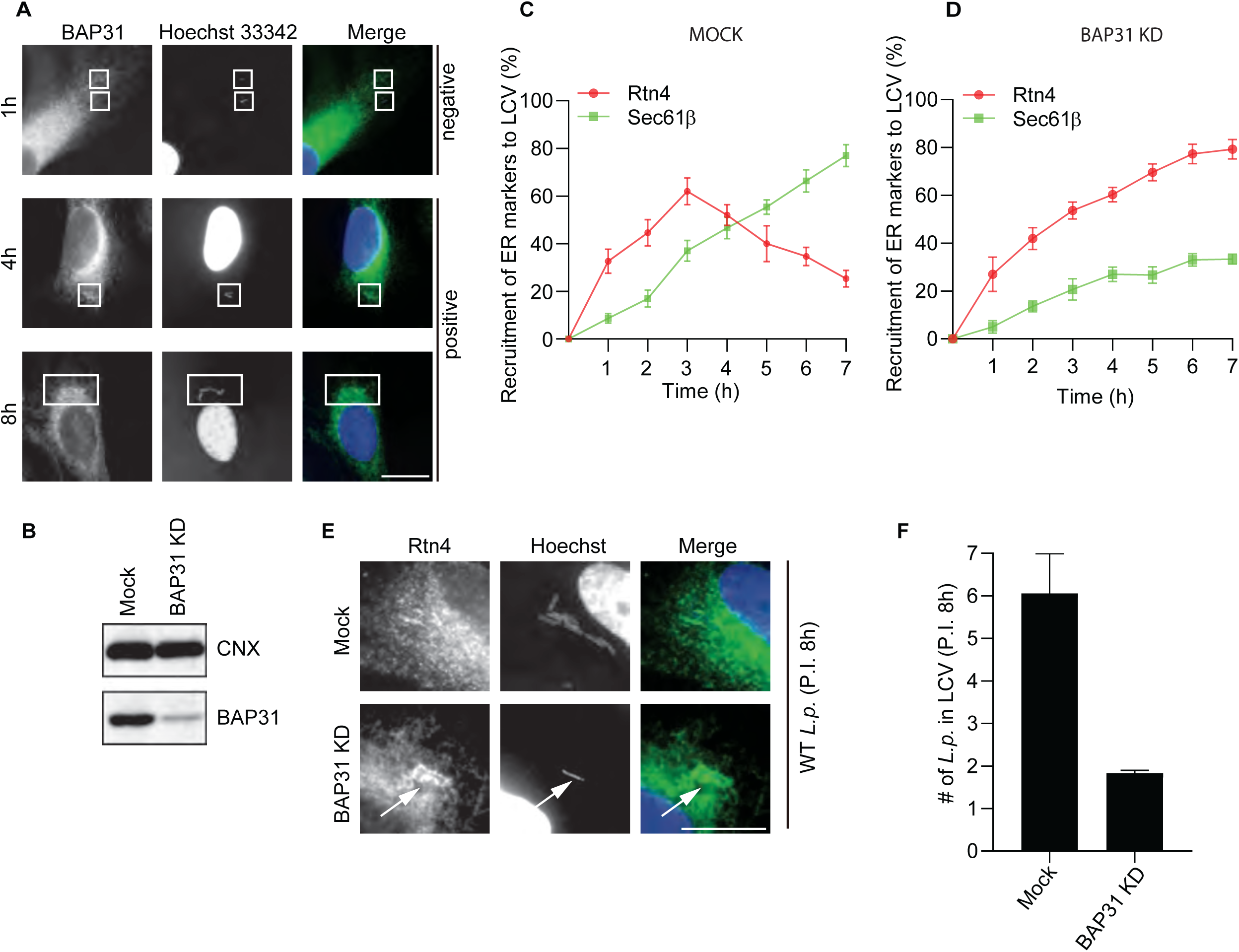
BAP31 is required for the sER to rER transition and efficiently replicate *Legionella.* (A) HeLa Fcγ cells were infected with *L.p*. for the indicated time, fixed, and stained with anti-BAP31 antibody and Hoechst 33342. White boxes indicate the LCV. (B) HeLa Fcγ cells were transfected with either mock or BAP31 siRNA for 72 hours and immunoblotted against the indicated antibodies. (C, D) Mock-treated, or BAP31 silenced HeLa Fcγ cells were infected with *L.p*. for the indicated time, fixed, and stained with anti-Rtn4 or -Sec61β antibodies and DAPI, and the vacuoles were quantified. Results were representative of three independent experiments. (100 vacuoles were scored in each experiment). Bars represent mean ± SD. (E) HeLa Fcγ cells were silenced with either mock or BAP31 for 72 hours, followed by infection with *L.p.* for 8 hours. Post-infection, cells were fixed and stained with anti-Rtn4 antibody and Hoechst for immunofluorescence analysis. Bar = 5 µm. (F) HeLa-Fcγ cells were silenced with either mock or BAP31 for 72 hours, followed by infection with *L.p.* for 8 hours. Post-infection, cells were stained with Hoechst, and the number of L.p. in the vacuoles was quantified. Results were representative of three independent experiments, and statistical significance was achieved using a student t-test (100 vacuoles were scored in each experiment). Bars represent mean ± SD.

### Lpg1152 brings BAP31 to LCVs and is required for the LCV maturation and replication of Legionella

*L.p* secretes over 330 effector proteins into the host cytosol; thus, finding the effector that binds to BAP31 is not trivial. We took advantage of *L.p.* strains that lack five large genomic island clusters and hence lack about ~70 effectors^36^ (**Fig. 5A**). Δ*P* is a pentuple strain lacking all five clusters (Δ*2ab*, Δ*3*, Δ*4a*, Δ*6a*, and Δ*7a*). To identify the effector(s) that are required for the recruitment of BAP31 on LCV, we analyzed WT, Δ*dotA* (which lacks the entire secretion system), Δ*P* (pentuple, which lacks all 5 clusters), and individual genomic cluster deletion strains (Δ*2ab*, Δ*3*, Δ*4a*, Δ*6a*, and Δ*7a*) of *L.p.* Excitingly; we found that BAP31 was not recruited on LCVs in the Δ*dotA*, Δ*P* and Δ*2ab* genomic strain, suggesting that one or more effector(s) present in the Δ*2a* region is required for the BAP31 recruitment on the LCVs **(Fig. 5B)**. No change was observed in the recruitment of BAP31 on LCVs from other cluster deletion strains, which includes Δ*3*, Δ*4a*, Δ*6a*, and Δ*7a* **(Fig. 5B)**, indicating that effectors from these regions are not essential for BAP31 recruitment. Further analysis of all the effectors on this island led to identifying a single effector, Lpg1152, that coimmunoprecipitated with BAP31 (**Fig. S4**). To confirm that Lpg1152 is an ER-localized protein, HeLa cells were transfected with 3xFLAG-Lpg1152, and immunofluorescence microscopy was performed with an anti-BAP31 antibody to detect endogenous BAP31. As expected, BAP31 colocalized with Lpg1152 **(Fig. 5C)**. Next, we set out to see if Lpg1152 interacts with BAP31 and BAP29. To this end, HEK293-Fcγ cells were transfected with either a 3X-FLAG vector (negative control) or 3x-FLAG-Lpg1152, followed by a co-IP using a FLAG tag antibody. Endogenous BAP31 and BAP29 were then immunoblotted. Surprisingly, only BAP31 but not BAP29 interacted with Lpg1152 **(Fig. 5D and 5E)**. To test if Lpg1152 drives BAP31 recruitment to the LCV, we generated a Δ*lpg1152* strain and a complemented Δ*lpg1152:*:3xFLAG-Lpg1152 strain. First, we confirmed that there is no significant difference in the infection efficiency of these mutant strains **(Fig. 5F)**. As expected, the Δ*lpg1152* strain showed a marked reduction in the recruitment of BAP31 to LCVs after 4 hours post-infection **(Fig. 5G)**. However, the recruitment of BAP31 to LCVs was restored with the Lpg1152 complemented strain (Δ*lpg1152::3xFLAG-Lpg1152*), suggesting a role for Lpg1152 in recruiting BAP31 to LCVs **(Fig. 5G)**. Surprisingly, we observed partial restoration of BAP31 recruitment to LCVs after 6 hours of Δ*lpg1152* infection, indicating that Lpg1152 is required for BAP31 recruitment early during infection. Perhaps other redundant effectors or Rab10 alone may be sufficient to recruit BAP31 later during infection. Next, we explored the role of Lpg1152 in recruiting sER and rER markers to LCVs. To this end, HeLa Fcγ cells were infected with WT, Δ*dotA*, Δ*lpg1152,* and Δ*lpg1152::3xFLAG-Lpg1152* strains, and cells were scored for the vacuoles positive with the sER and rER markers (Rtn4 and Sec61β). As expected, the Δ*lpg1152* strain showed enhanced recruitment of Rtn4 over Sec61β, suggesting a defect in transition to the rER 8 hours post-infection (**Fig. 5H**). In contrast, infection of HeLa Fcγ cells with Δ*lpg1152::*3xFLAG-Lpg1152 *L.p* strain restored the recruitment of Rtn4 and Sec61β to the WT *L.p* infection levels **(Fig. 5H)**. Finally, we elucidated the role of Lpg1152 in the replication of *Legionella* in HeLa Fcγ and U937 macrophage-like cells. We found that replication of the Δ*lpg1152 L.p.* strain was significantly inhibited in both cell types **(Fig. 5I and J)**. Notably, these phenotypes were rescued by Δ*lpg1152::3xFLAG-Lpg1152* WT strain, suggesting that Lpg1152 is required for optimal replication of *L.p.* Collectively, these results point towards a novel role for Lpg1152 in LCV maturation and replication.

**Figure 5.**
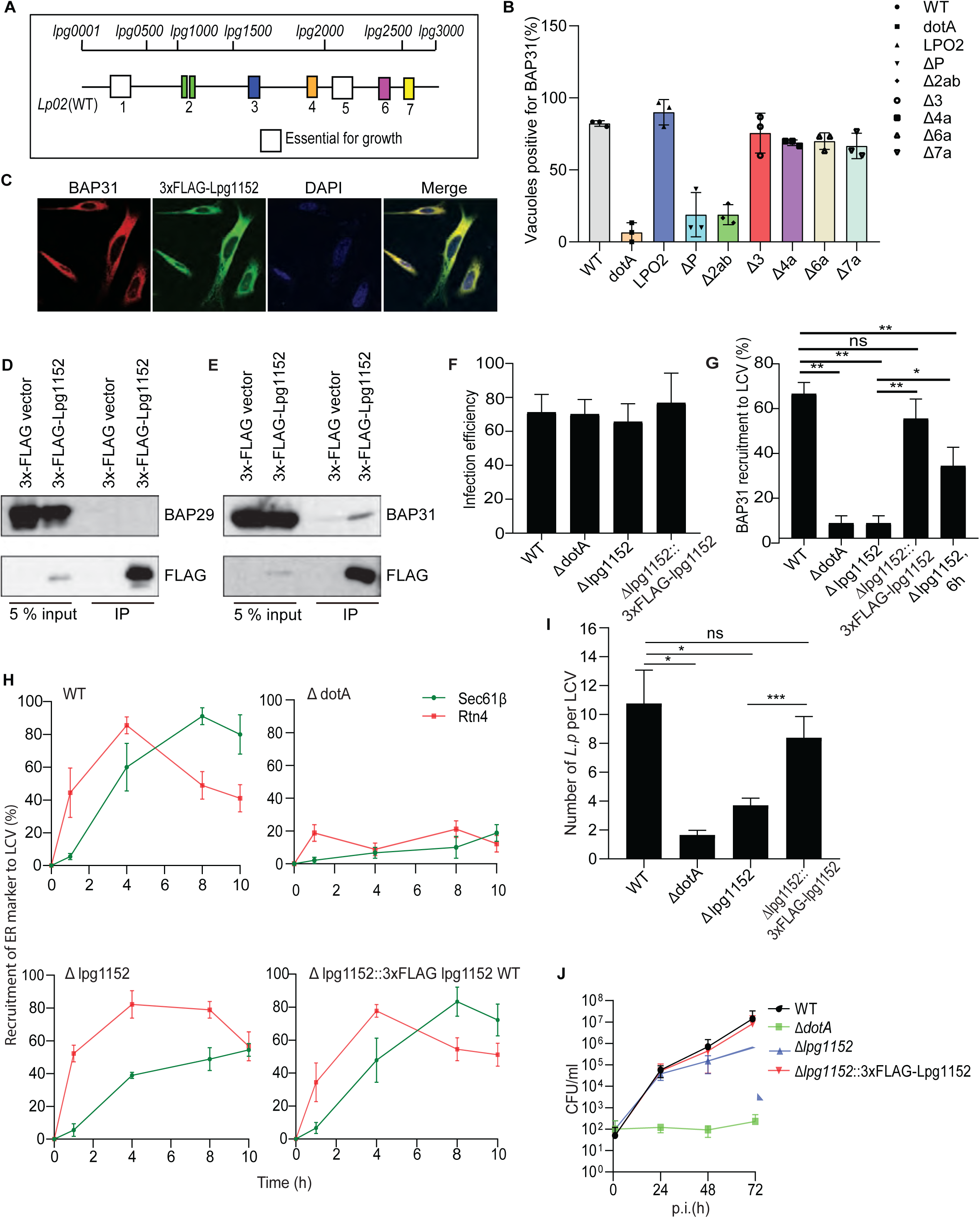
Lpg1152 brings BAP31 to LCVs and is required for optimal LCV maturation and replication of *Legionella.* **(A)** Schematic representation of the chromosomal organization of genomic islands of *L.p.* **(B)** HeLa Fcγ cells were infected with the indicated genomic island deletion mutant *L.p.* strains for 4 hours and processed for immunofluorescence against BAP31 antibody. The histogram represents the (%) of vacuoles positive for BAP31 post-infection with different *L.p.* strains. **(C)** HeLa Fcγ cells were transfected with 3xFLAG-Lpg1152 plasmid and processed for immunofluorescence microscopy using FLAG (green) and BAP31 (red) antibodies. Nuclei were stained with DAPI. (D, E) HEK293-Fcγ cells were transfected with the indicated plasmids. Cell extracts were subjected to immunoprecipitation assay post-transfection with anti-FLAG antibody and immunoblotted against the indicated antibodies. (F) HeLa Fcγ cells were infected with the indicated *L.p.* strains. Histograms represent the infection efficiency, i.e., the percentage of infected cells at MOI 10 1.5 hours after infection. (n=200 per triplicate experiment). (G) HeLa Fcγ cells were infected with the indicated strains of *L.p.* for the indicated time points. Following infection, LCV positive for BAP31 was scored using immunofluorescence analysis. (H) HeLa Fcγ cells were infected with the indicated strain, and the Rtn4 and Sec61β positive vacuoles were scored using immunofluorescence. Graphs represent the percentage recruitment of Rtn4 and Sec61β at the indicated time points post-infection. (I) HeLa-Fcγ cells were infected with the indicated *L.p* strains, and the number of *L.p.* per LCV was counted 8 hours after infection (n=30 per triplicate experiment). (J) U937 cells were infected with the indicated *L.p.* strains for 1 hour, and colony-forming units (CFU) were counted at the indicated time after infection. Data represent three independent experiments, and statistical significance was carried out with a Student t-test. Bar represents mean ± SEM.

### BAP29 plays an antagonistic role to Syntaxin 18 in maintaining sER and rER boundaries

We were intrigued by our finding that while BAP31 and BAP29 share 50% homology, *L.p.* effector Lpg1152 specifically interacted with BAP31 and not BAP29. Even though it was known that BAP31 and BAP29 form a complex^37^, the function of BAP29 remains undefined. Here, we uncovered a novel role of BAP proteins as Rab effectors that facilitate the transition of the LCV from the sER to the rER. Given that BAP31 shuttles between the sER and rER, but BAP29 does not, we wondered if BAP29 functions independently of BAP31. To address this, we first examined whether loss of BAP29 affects ER sub-compartment morphology. Typically, Sec61β, mainly seen on rER, is observed as a perinuclear sheet-like structure separated from the peripheral tube-like structure of sER (Rtn4). We observed this staining in mock or BAP31-silenced cells. However, when BAP29 is silenced, a significant distribution of Sec61β to peripheral tube-like structure that overlaps with Rtn4 was observed (white arrow, **Fig. 6A**), suggesting that compartmentation between sER and rER is collapsed when BAP29 is absent. Thus, BAP29 might play an independent role in maintaining rER/sER identity. To gain more critical insight into the BAP29 function, we next examined the effect of dysfunction of BAP29 on the patch-like structure of sER induced by silencing syntaxin 18 (Stx18)^38^. Stx18 was initially identified as a sER-localized soluble NSF attachment receptor (SNARE) and is known to maintain the organization of ER subdomains and ER exit sites^38^. Previous studies showed that the architecture of sER is drastically changed from tube-like to patch-like structure when Stx18 is depleted^38^. Consistent with this, we observed a similar patch-like structure of Rtn4 where it was separated from Sec61β when Stx18 was silenced (**Fig. 6B**). Given that this phenotype is thought to be caused by segregation of sER and rER, we hypothesized that additional depletion of BAP29 could restore the phenotype as it would promote mixing of sER with rER. To test this, we knocked down Stx18 alone (**Fig. 6C and 6D**) or in combination with BAP29 or BAP31 (**Fig. 6E**). In these conditions, the patch-like structure of sER marked by Rtn4 was observed in cells silencing Stx18 alone or both Stx18 and BAP31 (**Fig. 6E**). Notably, in cells where both BAP29 and Stx18 were depleted, the distribution pattern of Rtn4 was restored entirely from a patch-like to a tube-like structure (**Fig. 6E**), indicating that BAP29 plays a crucial role in the regulation of the boundary between sER and rER, and this function is independent of BAP31.

**Figure 6.**
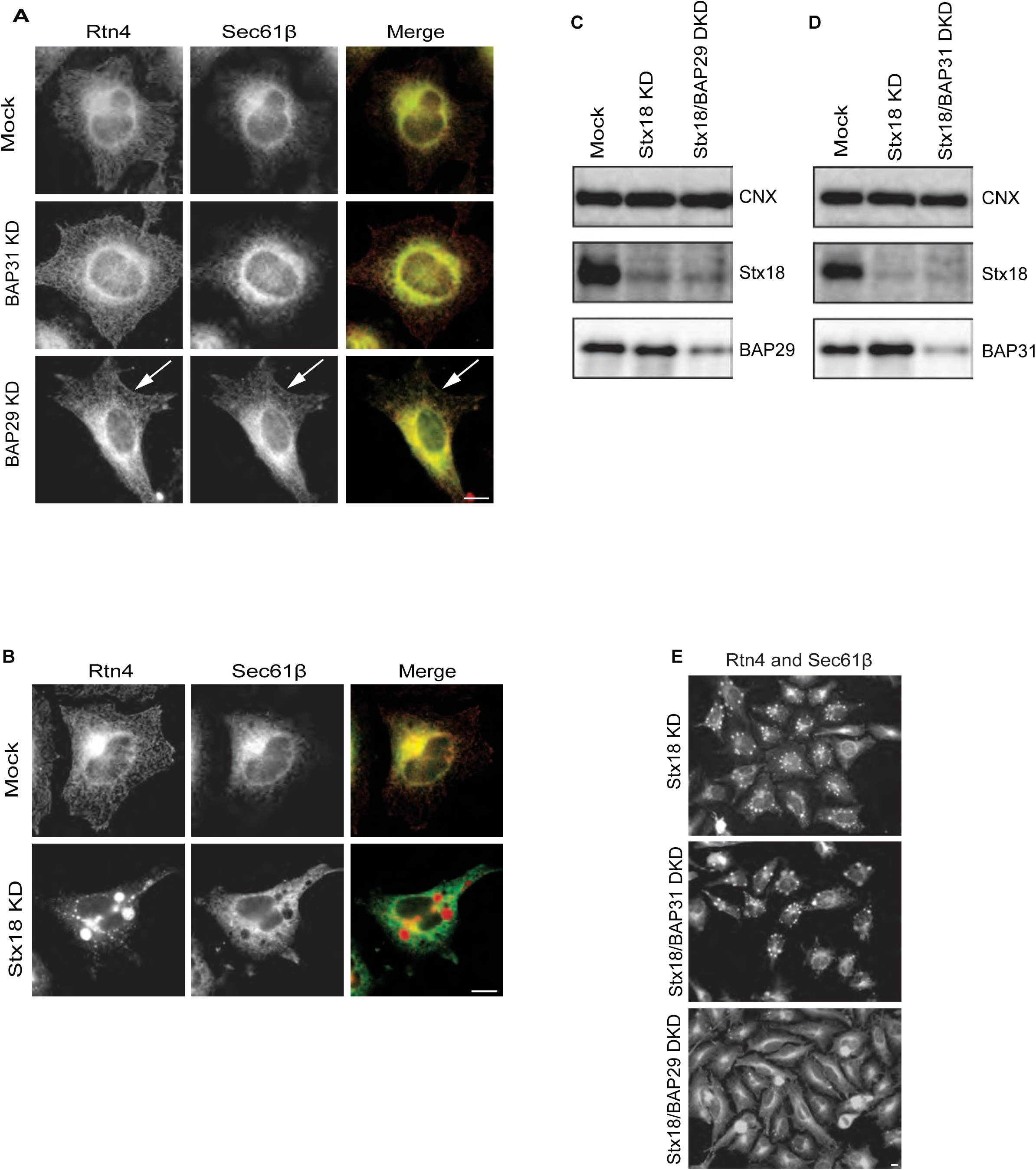
BAP29 is antagonistic to Syntaxin 18 function. (A) HeLa Fcγ cells were silenced with either mock, BAP31, or BAP29 siRNA for 72 hours. Post-silencing, cells were fixed and stained with anti-Rtn4 and Sec61β antibodies and were subjected to immunofluorescence imaging. Bar = 5 µm. White arrows indicate Sec61β localization to the cell periphery. (B) HeLa Fcγ cells were silenced with either mock or Stx-18 siRNA for 72 hours. Post-silencing, cells were fixed and stained with anti-Rtn4 and anti-Sec61β antibodies. Bar = 5 µm. (C, D) HeLa Fcγ cells were silenced for Stx18 alone or a combination of Stx18/BAP31 or Stx18/BAP29 siRNA. Following single or double silencing, cell extracts were prepared and immunoblotted against the indicated antibodies. (E) HeLa Fcγ cells were silenced for Stx18 alone, Stx18/BAP31, or Stx18/BAP29; following single or double silencing, cells were fixed and stained with an anti-Rtn4 antibody for immunofluorescence analysis. Bar = 5 µm.

## Discussion

Throughout history, pathogens have served as valuable tools for unraveling the intricacies of basic cell biology^39^. In recent years, *Legionella pneumophila* (*L.p*.) has emerged as particularly insightful in this regard^40^. While previous studies have indicated *L.p*.’s residence in an ER-like compartment, the localization of both smooth ER (sER) and rough ER (rER) markers within this compartment has presented conflicting interpretations. Moreover, the question remains whether this compartment is an integral part of the host ER or constitutes a distinct entity altogether. Though most pathogens enter the ER through retrograde trafficking, our use of FLIP and FRAP shows that *L.p.* does not reside within the host ER but establishes the *Legionella*-containing vacuole (LCV) by acquiring the ER membrane. To our knowledge, this is a unique manner of establishing an ER-like compartment and begs the question of what the benefits of creating a separate compartment are instead of utilizing the existing host compartment. This discovery provides a platform for future studies into the physical remodeling of ER membrane, which in turn could improve our understanding of processes such as ERphagy and stress-induced membrane remodeling.

Here, we describe a novel mechanism whereby *L.p.* diverges from the conventional retrograde trafficking route to co-opt the small GTPases Rab10 and Rab4, thus establishing an autonomous ER-like replicative niche, the LCV. We show that the knockdown of the small GTPase Rab10 prevents the LCV from acquiring ER identity, and the knockdown of Rab4 prevents the maturation of this compartment from sER to rER. Additionally, we elucidate that while this niche initially adopts an sER identity, *L.p.* orchestrates its maturation into rER through the actions of the proteins BAP31 and BAP29. These findings not only advance our understanding of *L.p.* pathogenesis but also shed light on fundamental aspects of cellular biology, highlighting the dynamic interplay between pathogens and host cells.

Furthermore, we demonstrate that the *L.p.* effector Lpg1152 binds to and recruits the host protein BAP31 to the LCV. While prior work showed that BAP31 shuttles between sER and rER^22^, our data establishes that it functions as a Rab effector to impart subcompartment identity. Indeed, knocking down BAP31 prevents the LCV from acquiring rER identity. We also show that the BAP31 homolog BAP29 resides on the LCV in a Rab10 and Rab4-dependent manner. While much less is known about BAP29 than BAP31, here, we discover a new role in maintaining sER and rER continuity in a syntaxin18-dependent manner.

Our findings elucidate new molecular players that define ER sub-compartment identity and facilitate the transition from sER to rER. This exciting finding further opens the possibility of exploring new mechanisms as to how a cell maintains its sER-rER ratio at any given time. Depending on the cell type, cells can have diverse protein and lipid synthesis needs. Thus, maintaining a correct sER-rER boundary can be vital for optimal cell function. Future work will help determine how separate compartmentalization benefits the pathogen and could lead to an improved understanding of the machinery and mechanisms involved in defining and remodeling ER subcompartments.

## MATERIAL AND METHODS

### Cell lines

HeLa, HeLa cells stably expressing the Fcγ receptor IIb (HeLa Fcγ cells) (a kind gift from the lab of Dr. Craig Roy at Yale University), HEK293-Fcγ and Hela Fcγ cells expressing GFP-Rtn4 and RFP-Sec1β were cultured in Dulbecco’s Modified Eagle’s Medium media (DMEM, GIBCO). U937 cells were cultured in RPMI-1640 (Corning) media. DMEM and RPMI-1640 media were supplemented with 10% heat-inactivated fetal bovine serum (FBS, VWR). U937 cells (a kind gift from Dr. Michael Bassik at Stanford University) were differentiated into macrophage-like cells using 20ng/ml phorbol 12-myristate 13 acetate (PMA, Sigma) for 72 hours, and cells were replated in media without PMA and allowed to recover for 72 hours before *Legionella pneumophila* (*L.p*) infection.

### Bacterial Strains and infection

*Lp01* WT and the isogenic Δ*dotA* strains are a kind gift from Dr. Craig Roy (Yale University). *Lp02* WT and the isogenic pentuple and individual genomic deletion cluster strains were kind gifts from Dr. Ralph Isberg (Tufts University). All *L.p.* strains were grown on charcoal yeast extract (CYE) agar plates supplemented with iron (FeNO_3_ 0.135g/10mL) and cysteine (0.4g/10mL). Chloramphenicol (10µg/mL), IPTG (0.1mM), and thymidine (100µg/mL) were added to CYE agar plates as needed. For infection experiments, primary patches of *L.p.* were grown for 2 days at 37°C on CYE plates. Single colonies of bacteria were picked and grown as heavy patches for 2 further days on CYE plates at 37°C. Heavy patches were then harvested, diluted in AYE broth supplemented with the required chemicals (as appropriate) and grown overnight at 37°C with shaking (220rpm) until the OD_600_ = ~3. HEK293 Fcγ cells were infected at a multiplicity of infection (MOI) of 50 or 5. For opsonization, *L.p.* bacteria were diluted in complete media and mixed with a *Legionella* polyclonal antibody (Invitrogen, Cat #PA1-7227) at 1:1,000 dilution. The mixtures were then incubated for 20 minutes at room temperature in a rotary mixer. Immediately after the addition of the opsonized *L.p.* mixtures, cells were centrifuged for 5 min at 1,000 rpm to facilitate spin-fection. Cells were then incubated at 37 °C for 60 minutes to allow bacterial uptake via the Fcγ receptor. After 60 minutes, the cells were washed with 1x PBS to remove extracellular bacteria and replenished with DMEM supplemented with 10% FBS. Infected cells were then cultured at 37 °C until time of harvest.

### Immunofluorescence

Hela Fcγ cells were seeded on 15mm glass coverslips in 12-well plates and transfected with indicated expression plasmids. 24 hours post-transfection, cells were washed with PBS and infected with *L.p.* at MOI = 10. One hour post-infection, cells were washed twice with PBS to remove unwanted extracellular bacteria. Following infection for the indicated times, cells were washed twice with PBS, fixed in 4% paraformaldehyde in PBS for 15 min at room temp. After fixation, extracellular bacteria were stained. Cells were then permeabilized with 0.2 % TritonX-100 in PBS, blocked in 2% bovine serum albumin (BSA) in PBS, and incubated with the indicated primary and secondary antibodies diluted in the blocking solution. Coverslips were mounted with ProLong Diamond Antifade Mountant with DAPI (Thermo Fisher Scientific) to stain host and bacterial nucleic acid.

### FLIP-FRAP

HeLa Fcγ cells stably expressing GFP-Rtn4 and RFP-Sec61β were infected with Halo-tagged WT *L.p.* for the indicated times. FRAP and FLIP techniques were used to photo bleach the sER marker Rtn4 (green) or the rER marker (red) Sec61β. For the FLIP experiment, the cell’s marked region (dotted squares) was photobleached every 10 seconds for 120 seconds, and timelapse images were taken every 1 second. For FRAP, the highlighted cell region was photobleached every 0.5 seconds for 40 seconds, and timelapse images were taken every 1-second post-bleaching.

### Proximity ligand assay (PLA)

Reagents of PLA were purchased from Sigma-Aldrich, and the assay was conducted according to the manufacturer’s protocol.

### CFU Assay

U-937 cells were stimulated with PMA (20ng/ml) for 72 hours and seeded in triplicates at 3×10^5^ cells/ well density. Post 72-hour PMA incubation, cells were washed with PBS and infected with WT, Δ*dotA*, Δ*lpg1152*, and Δ*lpg1152*::3X-FLAGLpg1152 strains of *L.p* for 1 hour (Day 0), 24 hour (Day 1), 48 hour (Day 2), and 72 hour (Day 3). One hour post-infection, cells were washed with PBS containing gentamycin (100mg/ml) to kill extracellular bacteria. After washing the cells with PBS, DMEM media containing gentamycin (100mg/ml) was added to wells for 45 minutes, and the cells were again washed three times with PBS. Post PBS washing, media was added back to wells, and cells were allowed to proceed for infection for the times mentioned earlier. Following infection, cells were lysed in 1 ml H_2_O and plated on CYE plates (with serial dilutions). Note: Only undiluted samples were used for the Δ*dotA* strain of *L.p.* since these mutants are defective in growth.

### Replicative Vacuole assay

HeLa Fcγ cells were transfected without or with siRNA against BAP31. At 72 hours after transfection, cells were infected with wild-type *L.p.* for eight hours and fixed, stained with anti-*L.p.* antibody for detection of extracellular *L.p.,* permeabilized with 0.2% TritonX-100, and re-stained with DAPI for detection of intracellular bacteria. The number of intracellular *L.p*. in each vacuole was scored.

### Immunoprecipitation

HEK293 Fcγ cells were seeded in 6-well plates and transfected with the indicated plasmid (3x-FLAG vector, 3x-FLAG-Rab4, 3x-FLAG-Rab5, 3x-FLAG-Rab10, or 3x-FLAG-Lpg1152). Post 24 hours, cells were washed with PBS and lysed in 300 µl of cell lysis buffer (150mM KCl, 25 mM Hepes-KOH pH7.4, 1% Triton-X100, protease inhibitor cocktail) on ice for 20 minutes. Lysates were spun down at 15,000 rpm for 10 minutes at 4°C, and detergent soluble supernatant fraction was collected and designated as cell lysate. Cell lysates were incubated with anti-FLAG M2 affinity beads (Sigma-Aldrich) for 1 hour at 4°C. After incubation, beads were washed three times with lysis buffer not containing the inhibitor cocktail, and the bound proteins were eluted by adding 20 µl of Laemmeli buffer and boiling for 5 min at 100°C.

### Western blotting

Sample-loaded SDS-gels were transferred onto the PVDF membrane (Merck-Millipore), and membranes were blocked in 5% milk or 2% BSA in TBS containing 0.1% Tween20 (TBS-T) for 1 hour at room temp. After blocking, membranes were incubated with primary antibodies for 1 hour at room temp or overnight at 4°C, washed with TBS-T three times, and incubated with horseradish peroxidase (HRP)-conjugated secondary antibodies (Bio-Rad) for 1 hour at room temp. The HRP signal was detected using an enhanced chemiluminescence reagent (Merck-Millipore).

### RNA interference

RNA duplexes used for targeting were Rab4 (5’-AACCTACAATGCGCTTACTAA-3’), Rab10 (5’-AAGAGTTGTACCTAAAGGAAA-3’), BAP29 (5’-AACTAAAAAGGATTTTGAAAA-3’), and BAP31 (5’-CAGCACTAAGCAAAAACTAGA-3’). RNA duplex used for targeting Stx18 was described previously. The RNA duplexes were purchased from Japan Bioservice, Inc. Transfection of the RNA duplexes was performed at a final concentration of 200 nM using Oligofectamine (Thermo Fisher Scientific) according to the manufacturer’s protocol.

### Subcellular fractionation

About 90% confluent HeLa Fcγ cells (three 15 cm dishes) were washed twice in PBS and then once in homogenization buffer (10 mM Hepes-KOH (pH7.4), 0.25 M sucrose, 1 mM EDTA, and protease inhibitor cocktail). After washing, cells were collected, suspended in 1 ml of homogenization buffer, and then homogenized by passaging 15 times through a 26G needle. The homogenate was centrifuged at *1,000 x g* for 10 min, and the resulting post-nuclear supernatant (PNS) was layered onto a 0-28% OptiPrep (Abbott Diagnostic Technologies) continuous gradients and centrifuged at *200,000 x g* for 6 hours. After centrifugation, 24 fractions (170 µl each) were collected from the top to bottom.

### Isolation of the smooth and rough ER fractions

Homogenate preparation from HeLa Fcγ cells (from five 15 cm dishes) was prepared as described above. After preparation of the PNS fraction, the PNS fraction was centrifuged at 10,000 x g for 10 min to remove the mitochondrial fraction. Then, the supernatant was additionally centrifuged at *100,000 x g* for 1 hour. The resulting pellet (total membrane fraction) was suspended in 1.5 ml homogenization buffer. The sucrose concentration of suspension was adjusted to 1.17 M by adding 2 M sucrose in homogenization buffer, and then 1.1 ml of suspension was overlaid on gradients of 1.1 ml of 1.15 M sucrose, 1.4 ml of 0.86 M sucrose, and 1.1 ml of 0.25 M sucrose in homogenization buffer. The gradients were centrifuged at *100,000 x g* for 3 hours, and then the two fractions, the smooth ER fraction at the upper portion of 1.17 M sucrose phase and the rough ER pellet were obtained. The membrane of the smooth ER fraction was collected by centrifugation at 100,000 x g for 1 hour, and the smooth ER pellet and the rough ER pellet were resuspended in a homogenization buffer.

## Supporting information

Supplemental Figures Low Res

## ACKNOWLEDGEMENTS

We thank Dr. Hana Kimura and Ady Steinbach for generating *L.p.* knockout (Δ*lpg1152*) and complemented strains (Δ*lpg1152:*:*3xFLAG-Lpg1152*) and conducting preliminary experiments with them. We thank Dr. Advait Subramanian and Tom Moss for critically reading and editing the manuscript. We thank Dr. Hana Kimura and the Center For Advanced Light Microscopy (CALM) for the FLIP and FRAP experiments. K.A. acknowledges Grant-in-Aid for scientific research from (grant nos 24790425, 26713016, 18H02656, and 20H05772), Uehara Memorial Foundation, Naito Foundation, and Takeda Science Foundation. S.M. acknowledges financial support from the National Institutes of Health (grant nos R01GM140440 and R01GM144378), the Pew Charitable Trust (grant no. A129837), a Bowes Biomedical Investigator award, and a gift fund from the Chan– Zuckerberg Biohub.

## AUTHOR CONTRIBUTIONS

K.A. and S.M. conceptualized the study and designed experiments. A.C., K.A., and S.M. wrote the manuscript. A.C., S.S., K.A., Y.Y., H.O., H.I., and Y.W. performed experiments. All authors analyzed data. K.A. and S.M. supervised the study. All authors reviewed and approved the manuscript.

