## Supplementary figures and images for "Legionella uses host Rab GTPases and BAP31 to create a unique ER niche"

### Supplemental Figures Low Res

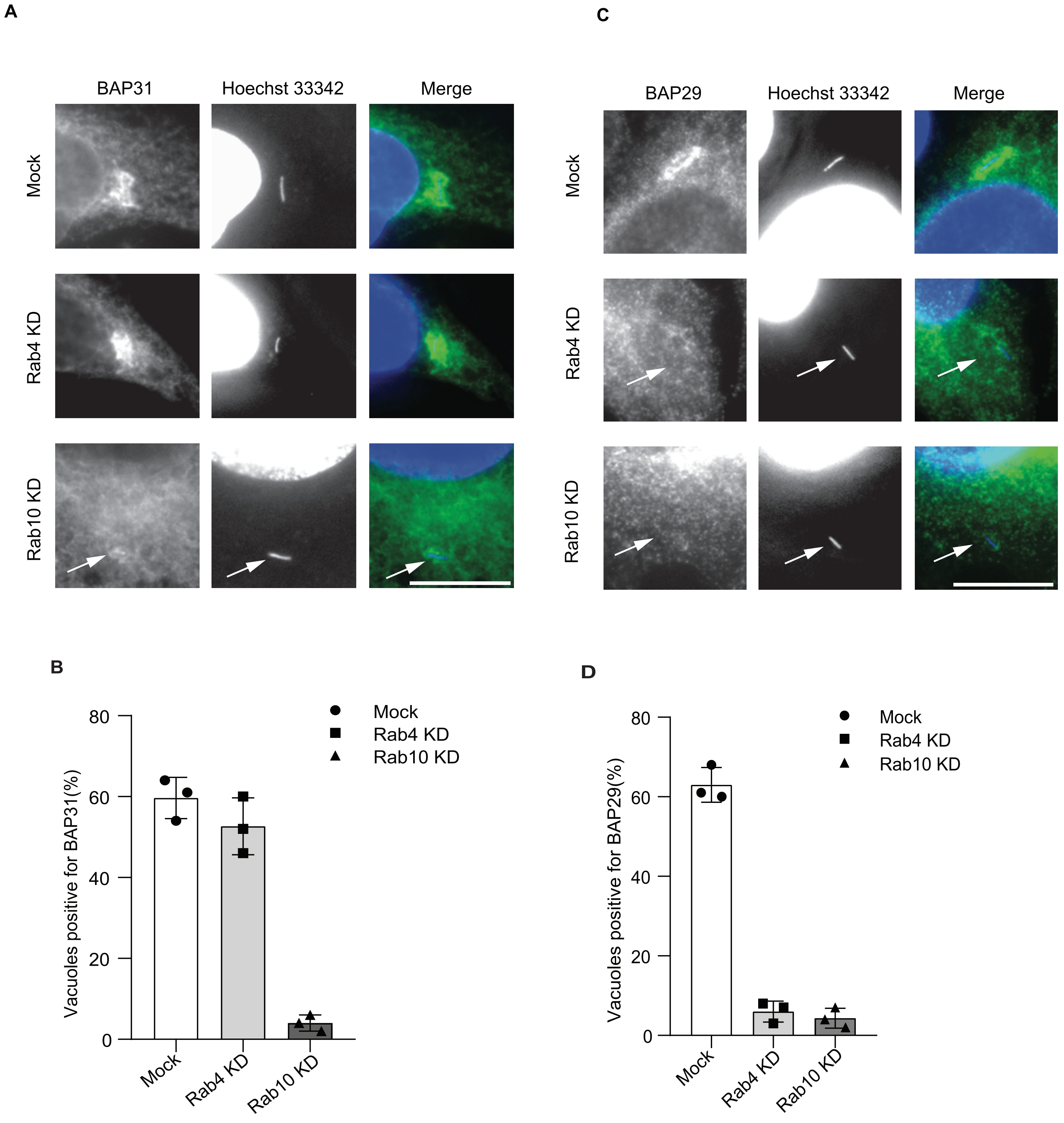

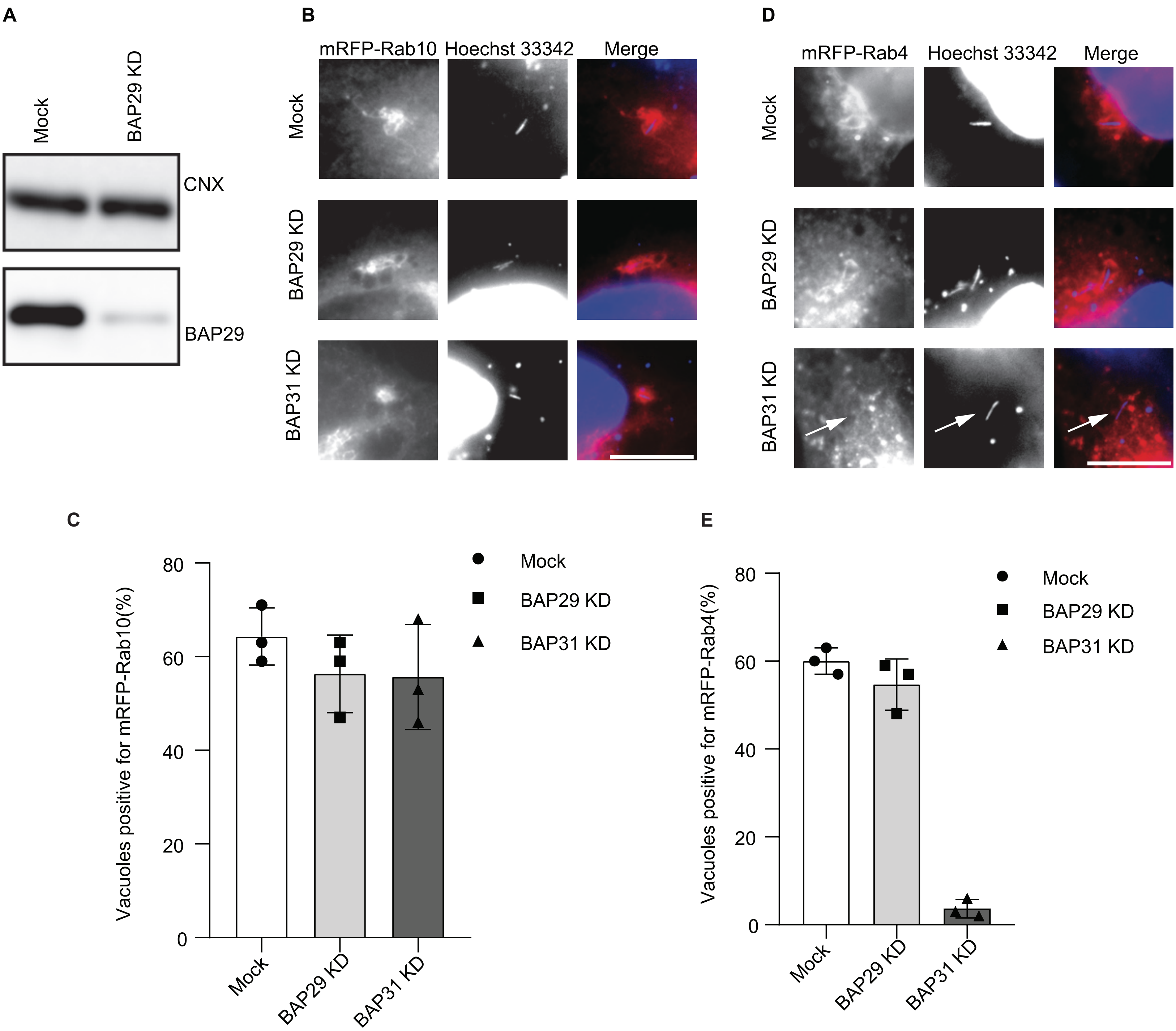

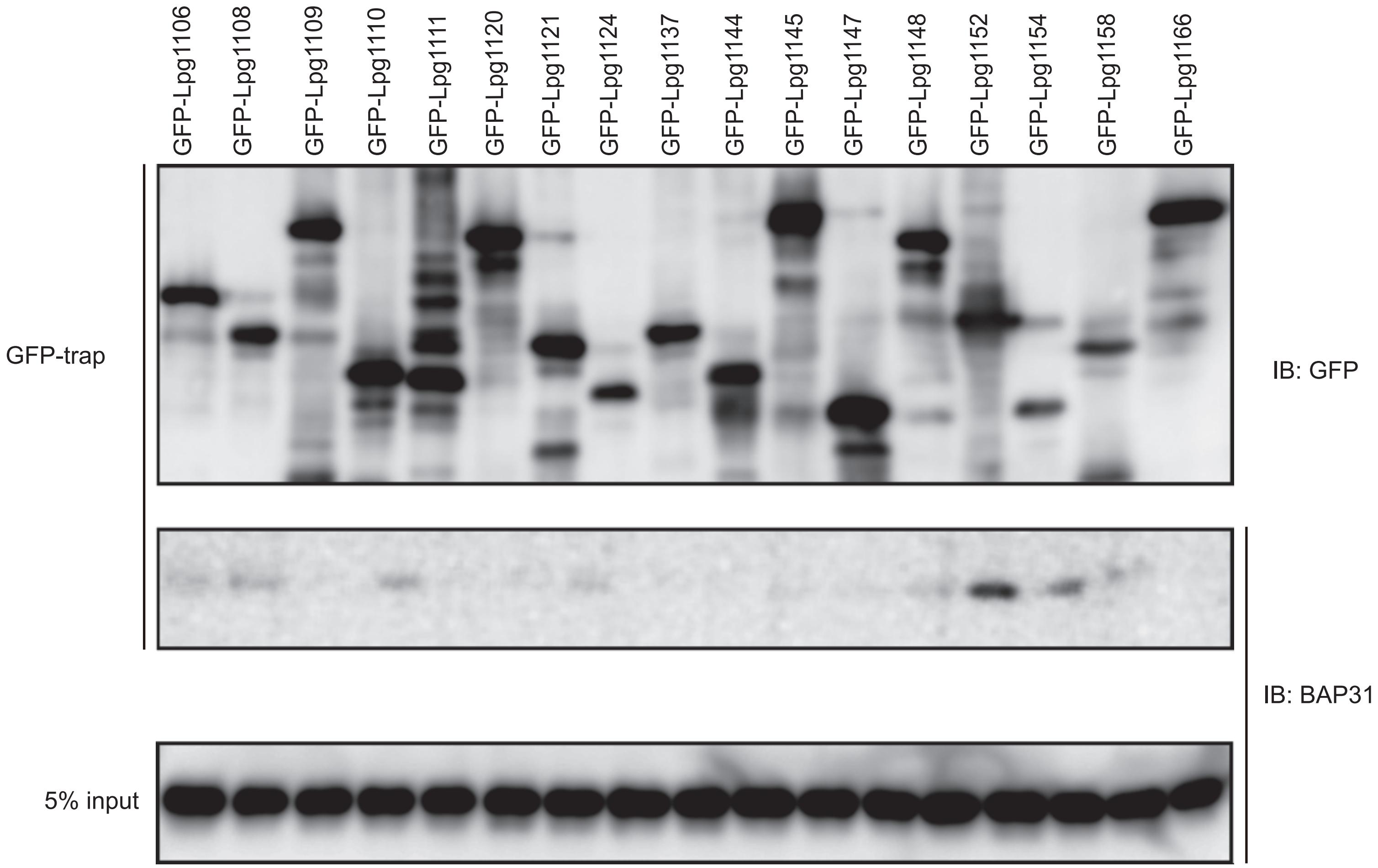
