## Supplemental Figure Legends for "Legionella uses host Rab GTPases and BAP31 to create a unique ER niche"

**Supplemental Figure Legend:**

**Figure S1.** **FLIP and FRAP show that LCV is associated with the s-ER early during infection but forms a separate r-ER niche at a later stage.** HeLa Fcγ cells stably expressing GFP-RTN4 and RFP-Sec61β were infected with Halo-tagged WT *L.p.* for the indicated time point, and time-lapse images were collected following either FLIP or FRAP photobleaching. (A) FLIP shows that GFP-Rtn4 is associated with the LCV at 3 hours post-infection, and after bleaching, the Rtn4 signal is lost, suggesting only an association, and not a separate compartment. (B and C) FLIP shows that RFP-Sec61β is associated with the LCV 6 hours post-infection, and after bleaching, the RFP-Sec61β signal around the LCV is not lost, suggesting that LCV is not contiguous with the host ER. (D, E) HeLa Fcγ cells stably expressing GFP-RTN4 and RFP-Sec61β were infected with Halo-tagged WT *L.p.* for 6 hours. RFP-Sec61β was photobleached around the LCV and the fluorescence was allowed to recover over time (FRAP). There is no recovery of fluorescence around the LCV suggesting that *L.p.* creates a separate LCV compartment independent of the host rER.

**Figure S2. Rab10 and Rab4 are required for BAP31 and BAP29 localization to the LCV.** (A) HeLa-Fcγ cells were silenced with either mock, Rab4, or Rab 10 siRNA for 72 hours, followed by infection with WT *L.p.* for 4 hours. Post-infection, cells were fixed and stained with anti-BAP31 antibody and Hoechst 33342 and processed for immunofluorescence imaging. Bar = 5 µm. (B) Quantitation of vacuoles positive for BAP31 following Rab4 and Rab 10 silencing. Data represent three independent experiments, and statistical significance was achieved with one-way ANOVA (100 vacuoles were scored in each experiment). Bars represent mean ± SD. (C) HeLa-Fcγ cells were silenced with either mock, Rab4, or Rab 10 siRNA for 72 hours, followed by infection with WT *L.p.* for 4 hours. Post-infection, cells were fixed and stained with anti-BAP29 antibody and Hoechst 33342 and processed for immunofluorescence imaging. Bar = 5 µm. (D) Quantitation of vacuoles positive for BAP29 following Rab4 and Rab 10 silencing. Data represent three independent experiments, and statistical significance was achieved with one-way ANOVA (100 vacuoles were scored in each experiment). Bars represent mean ± SD.

**Figure S3. The effect of BAP31 and BAP29 on recruitment of Rab4 or Rab10 on the LCV.** (A) HeLa Fcγ cells were transfected with either mock or BAP29 siRNA. Following transfection, cell extracts were subjected to immunoblotting against the indicated molecules. (B) HeLa-Fcγ cells expressing mRFP–Rab10 were silenced with either mock, BAP29, or BAP31 siRNA for 72 hours, followed by infection with WT *L.p.* for 4 hours. After infection, cells were fixed and stained with Hoechst 33342 and processed for immunofluorescence imaging. Bar = 5 µm. (C) Quantitation of vacuoles positive for mRFP-Rab10 following BAP29 and BAP31 silencing. Data represent three independent experiments, and statistical significance was achieved with one-way ANOVA (100 vacuoles were scored in each experiment). (D) HeLa-Fcγ cells expressing mRFP–Rab4 were silenced with either mock, BAP29, or BAP31 siRNA for 72 hours, followed by infection with WT *L.p.* for 4 hours. After infection, cells were fixed and stained with Hoechst 33342 and processed for immunofluorescence imaging. Bar = 5µm. (E) Quantitation of vacuoles positive for mRFP-Rab4 following BAP29 and BAP31 silencing. Data represent three independent experiments, and statistical significance was achieved with one-way ANOVA (100 vacuoles were scored in each experiment). Bars represent mean ± SD.

**Figure S4.** **Lpg1152 interacts with BAP31**. HEK293-Fcγ cells were transfected with plasmids encoding GFP-effectors encoded in island 2. After 24 hours of transfection, cell lysates were prepared and precipitated with GFP-trap beads. 5% input of lysates and precipitated proteins were analyzed with the indicated antibodies.
